# Optimizing mevalonate pathway for squalene production in *Yarrowia lipolytica*

**DOI:** 10.1101/2020.05.03.075259

**Authors:** Huan Liu, Fang Wang, Li Deng, Peng Xu

**Affiliations:** Department of Chemical, Biochemical and Environmental Engineering, University of Maryland Baltimore County, Baltimore, MD 21250; College of Life Science and Technology, Beijing University of Chemical Technology, Beijing, China

**Keywords:** Oleaginous yeast, metabolic engineering, mevalonate pathway, squalene, mannitol cycle

## Abstract

Squalene is the gateway molecule for triterpene-based natural products and steroids-based pharmaceuticals. As a super lubricant, it has been used widely in health care industry due to its skin compatibility and thermostability. Squalene is traditionally sourced from shark-hunting or oil plant extraction, which is cost-prohibitive and not sustainable. Reconstitution of squalene biosynthetic pathway in microbial hosts is considered as a promising alternative for cost-efficient and scalable synthesis of squalene. In this work, we reported the engineering of the oleaginous yeast, *Y. lipolytica*, as a potential host for squalene production. We systematically identified the bottleneck of the pathway and discovered that the native HMG-CoA reductase led to the highest squalene improvement. With the recycling of NADPH from the mannitol cycle, the engineered strain produced about 180.3 mg/l and 188.2 mg/L squalene from glucose or acetate minimal media, respectively. By optimizing the C/N ratio, controlling the media pH and mitigating the acetyl-CoA flux competition from lipogenesis, the engineered strain produced about 502.7 mg/L squalene in shake flaks, a 28-fold increase compared to the parental strain (17.2 mg/L). We also profiled the metabolic byproducts citric acid and mannitol level and observed that they are reincorporated into cell metabolism at the late stage of fermentation. This work may serve as a baseline to harness *Y. lipolytica* as an oleaginous cell factory for production of squalene or terpene-based chemicals.

## 1. Introduction

*Yarrowia lipolytica* is an industrial oleaginous yeast that has been extensively engineered to synthesize lipophilic compounds, including lipids (Qiao, Wasylenko, Zhou, Xu, & Stephanopoulos, 2017), oleochemicals (P. Xu, Qiao, Ahn, & Stephanopoulos, 2016), carotenoids (Gao et al., 2017; Macarena Larroude et al., 2018), terpenoids (Jin, Zhang, Song, & Cao, 2019) and aromatic polyketides (Lv, Marsafari, Koffas, Zhou, & Xu, 2019) *et al.* The lipogeneity of this yeast makes it a superior host to produce chemicals that are derived from acetyl-CoA, malonyl-CoA, HMG-CoA and NADPHs. The compartmentalization of oil droplets into lipid bodies provides a hydrophobic environment to sequestrate many lipid-related compounds and mitigate the toxicity issues associated with lipophilic membrane damages. In addition, the ease of genetic manipulation, substrate flexibility and robust growth present us tremendous opportunity to upgrade low-value renewable feedstocks to high-value compounds. It has also been recognized as a ‘generally regarded as safe’ (GRAS) organism (Groenewald et al., 2014) in the food and nutraceutical industry. A large collection of customized genetic toolboxes, including YaliBricks gene assembly (Wong, Engel, Jin, Holdridge, & Xu, 2017), CRISPR-Cas9 (Bae, Park, Kim, & Hahn, 2020; Macarena Larroude, Trabelsi, Nicaud, & Rossignol, 2020) or CRISPR-Cpf1 (Yang, Edwards, & Xu, 2020) genome editing, Cre-LoxP-based iterative chromosomal integrations (Lv, Edwards, Zhou, & Xu, 2019), transposon-based mutagenesis (Wagner, Williams, & Alper, 2018) and Golden-gate cloning (Celińska et al., 2017; Egermeier, Sauer, & Marx, 2019; M. Larroude et al., 2019), enabled us to rapidly modify its genome and evaluate many metabolic events to explore the catalytic diversity of this yeast beyond its regular portfolio of fatty acids, fatty alcohols, biofuels *et al*. Recent metabolic engineering effort in this yeast has allowed us to access more specialized secondary metabolites with pharmaceutical values, including sesquiterpenes (Marsafari & Xu, 2020), triterpenoids (Jin et al., 2019) and flavonoids (Lv, Marsafari, et al., 2019; Palmer, Miller, Nguyen, & Alper, 2020) *et al*.

Isoprenoids are a large group of natural products with diverse biological functions. An estimation of more than 70,000 isoprenoids, ranging from monoterpenes, sesquiterpenes, diterpenes and triterpenes have been discovered from nature (Moser & Pichler, 2019). Isoprenoids play a major role in maintaining membrane homeostasis, protein prenylation for subcellular targeting (Palsuledesai & Distefano, 2015), signal transduction, the deployment of plant defense pathways, and controlling the transcriptional activity of sterol-responsive-element-binding-proteins (SREBPs) (Shimano, 2001). The yeast-based mevalonate (MVA) pathway starts with acetyl-CoA condensation reactions, proceeds through the reduction of intermediate HMG-CoA *via* HMG-CoA reductase, which is the rate-limiting step and the molecular target to design many statins-related anti-cholesterol drugs (Xie & Tang, 2007). The universal five-carbon precursors isopentenyl diphosphate (IPP) and dimethylallyl diphosphate (DMAPP), derived from mevalonate, are condensed to make the farnesyl pyrophosphate (FPP), which later can be diversified to many sesquiterpenes or triterpenes. Squalene is a 30-carbon triterpene hydrocarbon synthesized from the condensation of two FPPs, which serve as the gateway molecule for all triterpenoids with tens of thousands of structural variations. Squalene possess strong antioxidant and anti-inflammatory activity and is widely used in the cosmetic industry as skin-compatible super-lubricant and hydration protectors (Spanova & Daum, 2011). Squalene emulsions were used as efficient adjuvants to enhance the immune response of certain vaccines (Spanova & Daum, 2011). Squalene is primarily sourced from shark liver, which poses significant ecological or ethical concerns related with shark-hunting. Reconstitution of squalene pathway in microbes may provide an alternative route to sustainably produce squalene from renewable feedstocks. A number of metabolic engineering studies have set the effort to engineer bacteria or bakers’ yeast to produce squalene, with improved yield and process efficiency. For example, a recent work identified that yeast peroxisome may serves as a dynamic depot to store squalene up to 350 mg/g dry cell weight (G.-S. Liu et al., 2020), despite the highly oxidative nature of peroxisome. In this work, we report the systematic optimization and engineering of the endogenous mevalonate pathway in *Yarrowia lipolytica* for efficient synthesis of squalene from simple synthetic media. We identified the bottlenecks of the mevalonate pathway and discovered alternative reducing equivalents (NADPH) pathways to improve squalene production. Our engineered strain produced up to 502.7 mg/L of squalene in shake flask. This work may set a foundation for us to explore oleaginous yeast as a chassis for cost-efficient production of squalene and triterpenoids in a long-term run.

## 2. Methods and Materials

### 2.1. Strains and culture conditions

*Escherichia coli* NEB5α high efficiency competent cells were obtained from NEB for plasmid construction, preparation, propagation and storage. The *Y. lipolytica* wild type strain W29 was purchased from ATCC (ATCC20460). The auxotrophic Po1g (Leu−) was obtained from Yeastern Biotech Company (Taipei, Taiwan). All strains and plasmids are listed in supplementary Table S2.

LB broth or agar plate with 100 µg/mL ampicillin was used to cultivate *E. coli* strains. Yeast rich medium (YPD) was prepared with 20 g/L Bacto peptone (Difco), 10 g/L yeast extract (Difco), and 20 g/L glucose (Sigma-Aldrich), and supplemented with 15 g/L Bacto agar (Difco) for solid plates. YNB seed medium was made with 1.7 g/L yeast nitrogen base (without amino acids and ammonium sulfate) (Difco), 5 g/L ammonium sulfate (Sigma-Aldrich), 0.69 g/L CSM-Leu (Sunrise Science Products, Inc.) and 20 g/L glucose. Selective YNB plates were made with YNB media supplemented with 15 g/L Bacto agar (Difco). In fermentation process, YNB fermentation medium with glucose as substrate and carbon/nitrogen molar ratio of 80:1 was made with 1.7 g/L yeast nitrogen base (without amino acids and ammonium sulfate) (Difco), 1.1 g/L ammonium sulfate (Sigma-Aldrich), 0.69 g/L CSM-Leu (Sunrise Science Products, Inc.) and 40 g/L glucose. YNB fermentation medium with sodium acetate as substrate and carbon/nitrogen molar ratio 80:1 was made with 1.7 g/L yeast nitrogen base (without amino acids and ammonium sulfate) (Difco), 0.825 g/L ammonium sulfate (Sigma-Aldrich), 0.69 g/L CSM-Leu (Sunrise Science Products, Inc.), 41 g/L sodium acetate. Glacial acetic acids were purchased from Sigma-Aldrich.

Phosphoric buffer solution (PBS) with pH 6.0 was made with 0.2 M Na_2_HPO4 and 0.2 M Na_2_HPO_4_, which was used to replace water to make YNB-glucose-PBS fermentation medium. Bromocresol purple was a pH-sensitive indicator which could change its color with the pH from 5.2-7.0 (El-Ashgar et al., 2012) and 40 mg/L bromocresol purple was added into fermentation medium to indicate pH variation. The pH of medium was regulated to 6.0 by 6. 0 M HCl in the fermentation process. The components in glucose-YNB media with C/N ratio 60:1, 40:1, 20:1, 10:1 were as same as them in C/N ratio 80:1 except the content of ammonium sulfate changed to 1.47 g/L, 2.2 g/L, 4.4 g/L, 8.8 g/L respectively, to explore the effect of C/N ratio on squalenen accumulation. And 1 mg/L cerulenin solution prepared with dimethylsulfoxide (DMSO) was added into fermentation medium to inhibit precursor (fatty acids biosynthesis) competing pathway.

### 2.2 Genetic transformation of *Y. lipolytica*

All plasmids constructed were transformed into the *Y. lipolytica* host strain Po1g ΔLeu using the lithium acetate/single-strand carrier DNA/PEG method (Chen, Beckerich, & Gaillardin, 1997). And single fresh *Y. lipolytica* colonies were picked from YNB selective plates and inoculated into YNB seed media, which were grown at 30 °C for 48 h. For tube test, 100 μL seed cultures were inoculated into 5 mL fermentation media in 50 mL tube. 600 μL seed cultures were inoculated into 30 mL fermentation media in 250 mL shake flasks with 250 rpm and 30 °C. Time series samples were taken for analyzing biomass, sugar content, and squalene titer.

### 2.3. Analytical methods

Four OD units of liquid yeast cell was harvested and subsequently was pelleted by centrifugation at 14,000 rpm for 5min. Water was completely withdrawn and 500 uL 0.5 M sodium methoxide (sodium hydroxide dissolved in pure methanol) was used to resuspend the cell pellet. The mixture was kept at room temperature with shaking for 2 hours at 1,200 rpm with a high-duty vortex to allow complete saponification of lipids and extraction of squalene. Then 400 uL hexane was added to extract squalene. The mixture was vortexed at room temperature for 10 min and hexane phase was directly injected to gas chromatography-FID (GC-FID) for squalene analysis. Gas chromatography–flame ionization detector (GC-FID) system (Agilent 7820A) equipped with HP-5 column (30 m × 320 μm × 0.25 μm) was used to detect squalene, using helium as the carrier gas with a linear velocity of 2 ml/min. The column temperature profile was 175 ℃ for 3 min, 20 ℃/min ramping to 200 ℃ and holding for 3 min, and then 20 ℃/min ramping to 260 ℃ and holding for 4 min.

The cell growth was monitored by measuring the optical density at 600 nm (OD600) with a UV-vis spectrophotometer that could also be converted to dry cell weight (DCW) according to the calibration curve DCW: OD600 = 0.33:1 (g/L). The fermentation broth was centrifuged at 14,000 rpm for 5 min and the supernatant was used for analyzing the concentration of glucose, mannitol, and acetic acid by HPLC with a refractive index detector and Supelcogel TM Carbohydrate column. The column was eluted with 10 mM H_2_SO_4_ at a column temperature of 50 ℃, a refractive index detector temperature of 50 ℃, and a flow rate of 0.4 mL/min.

### 2.4. Plasmid and pathway construction

All primers are listed in supplementary Table S1. All restriction enzymes were purchased from Fisher FastDigest enzymes. Plasmid miniprep, PCR clean-up, and gel DNA recoveries were using Zyppy and Zymoclean kits purchased from Zymo research. All the genes were PCR-amplified with the primers from genomic DNA of *Y. lipolytica*, *S. cerevisiae*, *E. coli*, *B. subtilis, Aspergillus nidulans*, respectively (Supplymentary Table S1 and Table S2). All these genes were inserted into downstream of the *Y. lipolytica* TEF-intron promoter in the pYLXP’ vector backbone (Wong et al., 2017) at the SnaBI and KpnI site *via* Gibson assembly (Gibson et al., 2009). Upon sequence verification by Genewiz, the restriction enzyme *Avr*II, *Nhe*I, *Not*I, *Cla*I and *Sal*I (Fermentas, Thermo Fisher Scientific) were used to digest these vectors, and the donor DNA fragments were gel purified and assembled into the recipient vector containing previous pathway components in compliance with the YaliBricks subcloning protocol (Wong et al., 2017; Wong, Holdridge, Engel, & Xu, 2019). All assembled plasmids were verified by gel digestion and were subsequently transformed into the *Y. lipolytica* host strain Po1g ΔLeu using the lithium acetate/single-strand carrier DNA/PEG method (Chen et al., 1997). In chromosomal integration process, pYLXP’ vector assembled with functional genes was linearized by restriction enzyme NotI (Fermentas, Thermo Fisher Scientific). The linear fragment was transformed into the *Y. lipolytica* host strain Po1g ΔLeu and cultivated on CSM-Leu minimal media (agar plate) for colony screening. The screened colony was later cultivated in YPD media and genomic DNA was extracted with Wizard genomic kits (Promega). Then the genomic samples were used as template for PCR verification of the integrated gene with gene-specific primers.

## 3. Results and discussions

### 3.1 Debottlenecking mevalonate pathway for squalene production

In yeast, squalene was primarily synthesized from the mevalonate (MVA) pathway (Fig. 1). Staring with acetyl-CoA condensation, yeast uses a number of critical enzymes to synthesize squalene, including acetoacetyl-CoA thiolase (Erg 10, YALI0B08536g), HMG-CoA reductase (YALI0E04807g), mevalonate kinase (Erg 12, YALI0B16038g), phosphomevalonate kinase (Erg 8, YALI0E06193g), mevalonate pyrophosphate decarboxylase (MVD1, YALI0F05632g), farnesyl pyrophosphate synthase (Erg20, YALI0E05753g), geranyl pyrophosphate synthase (YALI0D17050g) and squalene synthase (SQS1, YALI0A10076g). Genome annotation indicates that *Y. lipolytica* contains the complete mevalonate pathway (Fig. 1). In MVA pathway, HMG-CoA reductase was reported as the rate-limiting metabolic step in squalene accumulation (RODWELL, NORDSTROM, & MITSCHELEN, 1976). In addition, there was almost no squalene accumulated by native *Y. lipolytica* due to the quick consumption of squalene by downstream ergosterol synthase. After we overexpressed the endogenous squalene synthase gene (*SQS*), squalene production was increased to 17.25 mg/L at 120 h with chemically-defined complete synthetic media (CSM-leu) in test tube. With this as a starting strain, we investigated the effect of three HMG-CoA reductases (encoded by HMG) on squalene production. The three HMGs were derived from *Saccharomyces cerevisiae*, *Silicibacter pomeroyi* and *Y. lipolytica*. Truncated form of HMG-CoA reductase devoid of N-terminal membrane targeting signal has been proven to be effective in improving isoprenoid production in *Saccharomyces cerevisiae* (encoded by *SctHMG*) (Polakowski, Stahl, & Lang, 1998; Thompson, Kwak, & Jin, 2014). When co-expressed with endogenous SQS, the strain with the truncated HMG1 (SctHMG) led to squalene production at 83.76 mg/L (Fig. 2A), indicating that overexpression of HMG-CoA reductase was beneficial for squalene production. To test whether other sources of HMG-CoA reductase could display better functions, we co-expressed *SpHMG* from *Silicibacter pomeroyi* and endogenous *ylHMG* with SQS, respectively. A low yield of squalene (9.24 mg/L) was produced in the strain expressing *SpHMG*. This result was consistent with previous findings that HMG from *Silicibacter pomeroyi* was highly specific for NADH (Meadows et al., 2016) and this bacterial-derived enzymes could not be directly translated to yeast system. When endogenous *ylHMG* was co-expressed with SQS (strain *HLYaliS01*), the engineered strain yielded 121.31 mg/L squalene at 120 h in test tube, demonstrating the potential of using *Y. lipolytica* as a platform to synthesize various terpenes. We also tested the truncated form of *ylHMG* sequence (YALI0E04807p), of which the first 495 nucleotides that encode the 165 amino acid N-terminal domain responsible for membrane localization (ER targeting) were removed. The remaining C-terminal residues containing the catalytic domain and an NADPH-binding region (Gao et al., 2017) were overexpressed. We then overexpressed the truncated *ylHMG* (t495*ylHMG*) to compare how the variation of *ylHMG* may improve squalene synthesis. Contrary to our hypothesis, removal of the N-terminal 495bp of *ylHMG* exhibits adverse effect on both squalene production and cell growth (Fig. 2A), indicating that the N-terminal membrane-binding domain plays a critical role in squalene synthesis.

**Fig. 1.**
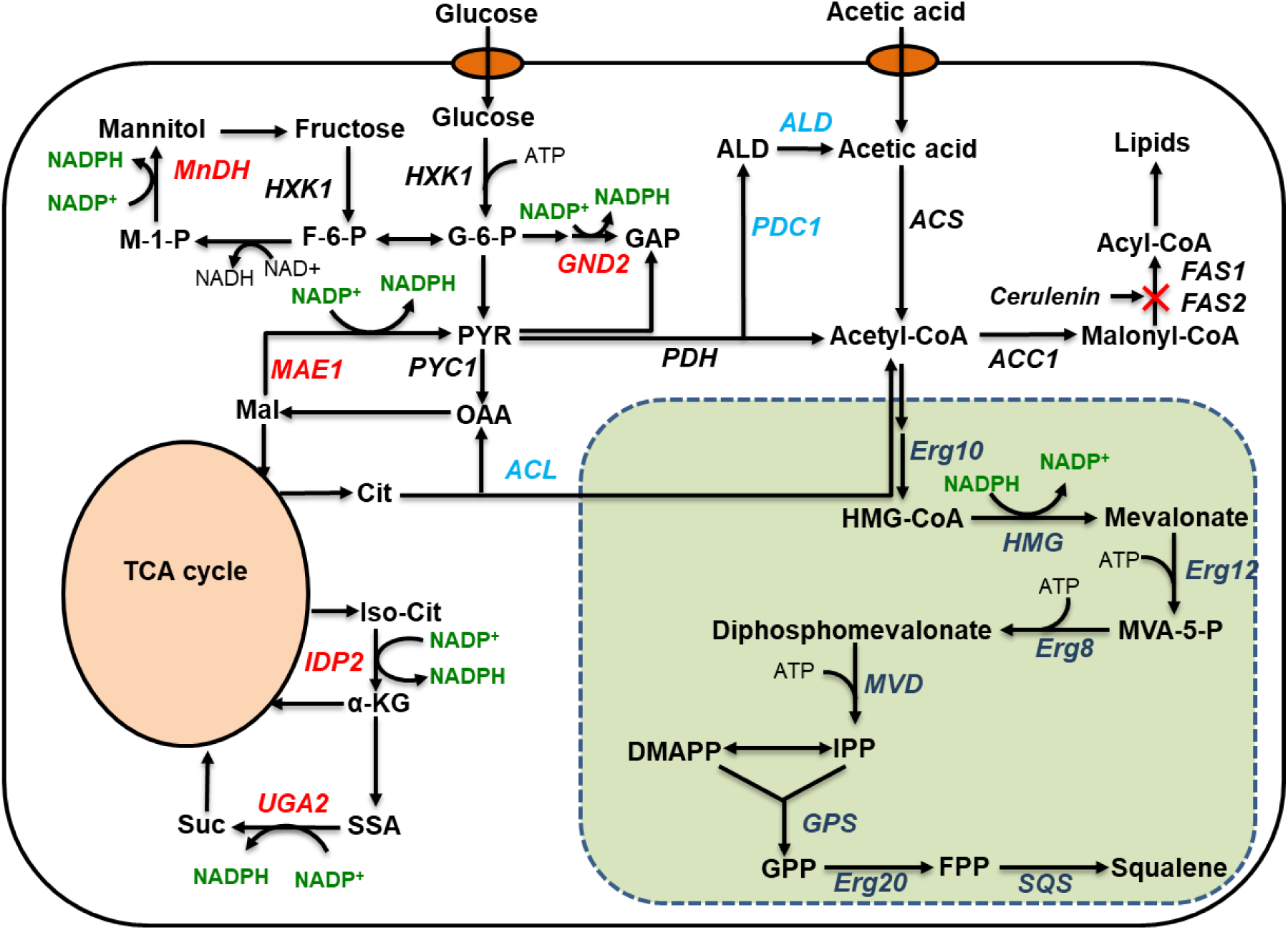
Metabolic pathway for squalene synthesis in oleaginous yeast. MnDH, mannitol dehydrogenase; HXK, hexokinase; MAE1, malic enzyme; ACL2, ATP citrate lyase; IDP2, cytosolic NADP-specific isocitrate dehydrogenase; UGA2, succinate semialdehyde dehydrogenase; PYC1, pyruvate carboxylase; PDC1, pyruvate decarboxylase; ALD, aldehyde dehydrogenase; PDH, pyruvate dehydrogenase complex; ACS, acetyl-CoA synthase; FAS1 and FAS2, fatty acid synthase; ACC1, acetyl-CoA carboxylase; HMG, HMG-CoA reductase; Erg 10, acetoacetyl-CoA thiolase; Erg 12, mevalonate kinase; Erg 8, phosphomevalonate kinase; MVD, mevalonate pyrophosphate decarboxylase; Erg 20, farnesyl pyrophosphate synthetase; GPS, geranyl pyrophosphate synthase; SQS, squalene synthase.

**Fig. 2.**
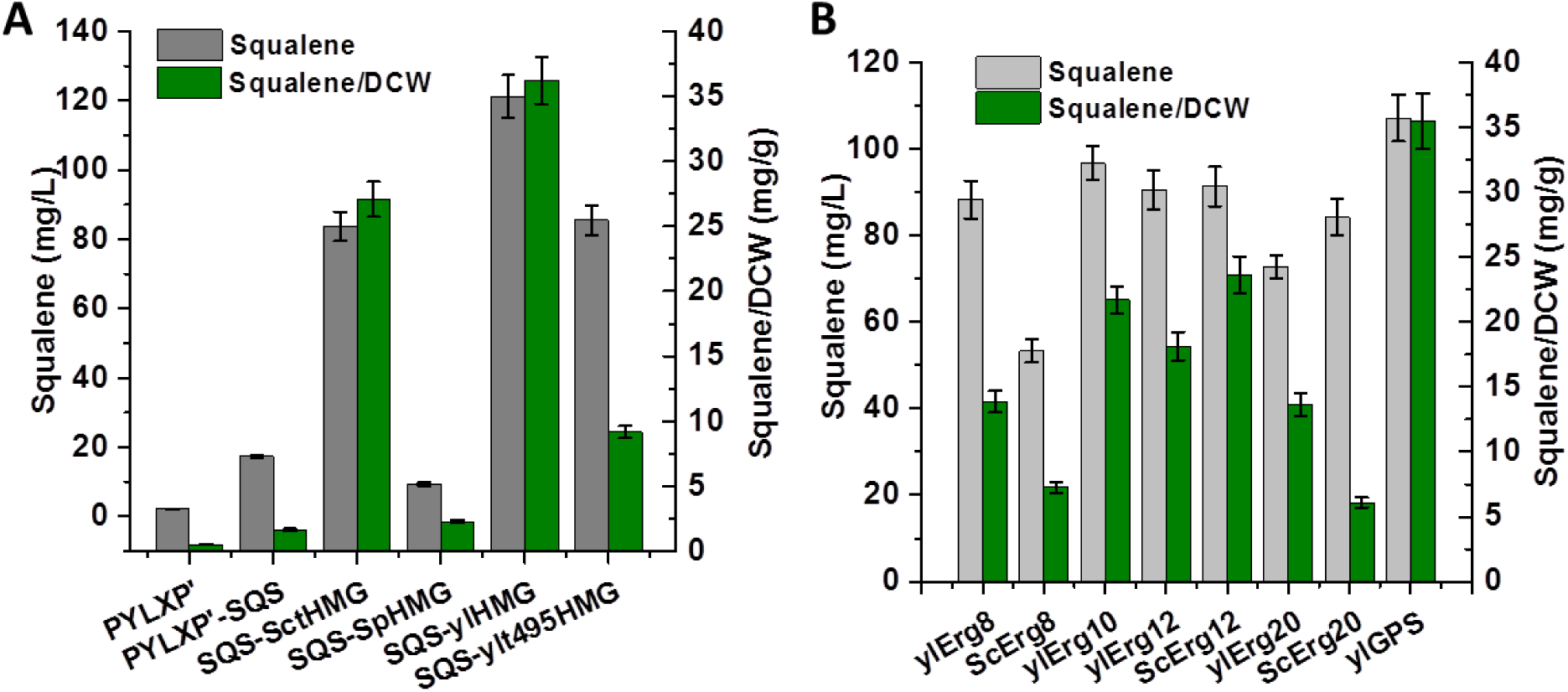
Comparison of different HMG-CoA reductase and identification of rate-limiting steps of endogenous mevalonate pathway in *Y. lipolytica*.

In addition to the overexpression of the endogenous SQS and *ylHMG* genes, we also tested whether the expression of other genes in the MVA pathway would improve squalene production, including ylErg8 encoding phosphomevalonate kinase, ylErg10 encoding acetoacetyl-CoA thiolase, Erg12 encoding mevalonate kinase, and ylErg20 encoding farnesyl pyrophosphate synthetase; ylGPS encoding geranyl pyrophosphate synthase. And ylErg8, ylErg10, Erg12 and ylErg20 from *S. cerevisiae* were also overexpressed to compare how the variation of these genes may enhance squalene synthesis. As shown in Fig. 2B, co-overexpression of ylErg8, ylErg10, Erg12 could not further improve squalene synthesis, regardless of the source of the gene. Among all of these combinations (Fig. 2B), the highest squalene production was obtained for the strain in which ylGPS and SQS-ylHMG1 were overexpressed, with titer of 107.08 mg/L and a specific production of 36.24 mg/g DCW, which is still lower than the strain only expressing SQS-ylHMG1. These results indicate that sequential overexpression of the genes involved in the MVA pathway could not further improve the carbon flux toward squalene, possibly due to the stringent regulation of MVA pathway at multiple nodes, including ergosterol-mediated feedback inhibition or SREBP-related transcriptional repression.

### 3.2 Augmenting NAPDH and acetyl-CoA precursor pathways to improve squalene production

NADPH as the primary biological reducing equivalent protects cell from oxidative stress and extend carbon-carbon backbones, which was also reported as the major rate-limiting precursor in fatty acids synthesis in oleaginous species (Qiao et al., 2017; Wasylenko, Ahn, & Stephanopoulos, 2015). HMG-CoA reductase (HMGR) is the first rate-limiting enzyme in the mevalonate pathway and plays critical role in regulating squalene biosynthesis (Ma et al., 2019). 3-hydroxy-3-methylglutaryl coenzyme A (HMG-CoA) is reductively hydrolyzed to mevalonate by releasing coenzyme A with NADPH as reducing equivalent (Cao, Wei, Lin, & Hua, 2017). Based on previous work, source of cytosolic NADPH in the Baker’s yeast may originate from various alternative routes depending on the carbon source and genetic background of the yeast strain (Huan Liu, Marsafari, Deng, & Xu, 2019; Minard, Jennings, Loftus, Xuan, & McAlister-Henn, 1998; Minard & McAlister-Henn, 2005). With glucose as carbon sources, cytosolic NADPH primarily relies on the pentose phosphate pathway. Other cytosolic NADPH pathways include NADP-specific isocitrate dehydrogenase (IDP2), malic enzyme (ylMAE), mannitol dehydrogenase (ylMnDH1, ylMnDH2), 6-phosphogluconate dehydrogenase (ylGND2) and succinate semialdehyde dehydrogenase (ylUGA2) (Huan Liu, Monireh Marsafari, Li Deng, et al., 2019) (Fig. 1). In this work, we tested a collection of auxiliary cytosolic NADPH pathways and investigated how these pathways may enhance squalene production and cellular fitness on the basis of co-expression *SQS-ylHMG* (Fig. 3A). Among these chosen NADPHs, mannitol dehydrogenase (ylMnDH2, encoded by YALI0D18964g) presented the best results to improve squalene production. Mannitol, a more reduced sugar alcohol compared to glucose, played an essential role in modulating cytosolic NADPHs through the mannitol cycle. This could partially explain why mannitol was the major byproduct during lipid accumulation phase in *Y. lipolytica* (P. Xu, Qiao, & Stephanopoulos, 2017). When ylMnDH2 was overexpressed with SQS and ylHMG (*strain HLYaliS02*, Supplymentary Table S2), the engineered strain produced 11% more squalene with volumetric production titer increased to 135.22 mg/L, despite relatively decreased yield of 32.33 mg/g DCW (Fig. 3A). This is possibly ascribed to the increased cell fitness and lipid content after enhancing the supplement of NADPH.

**Fig. 3.**
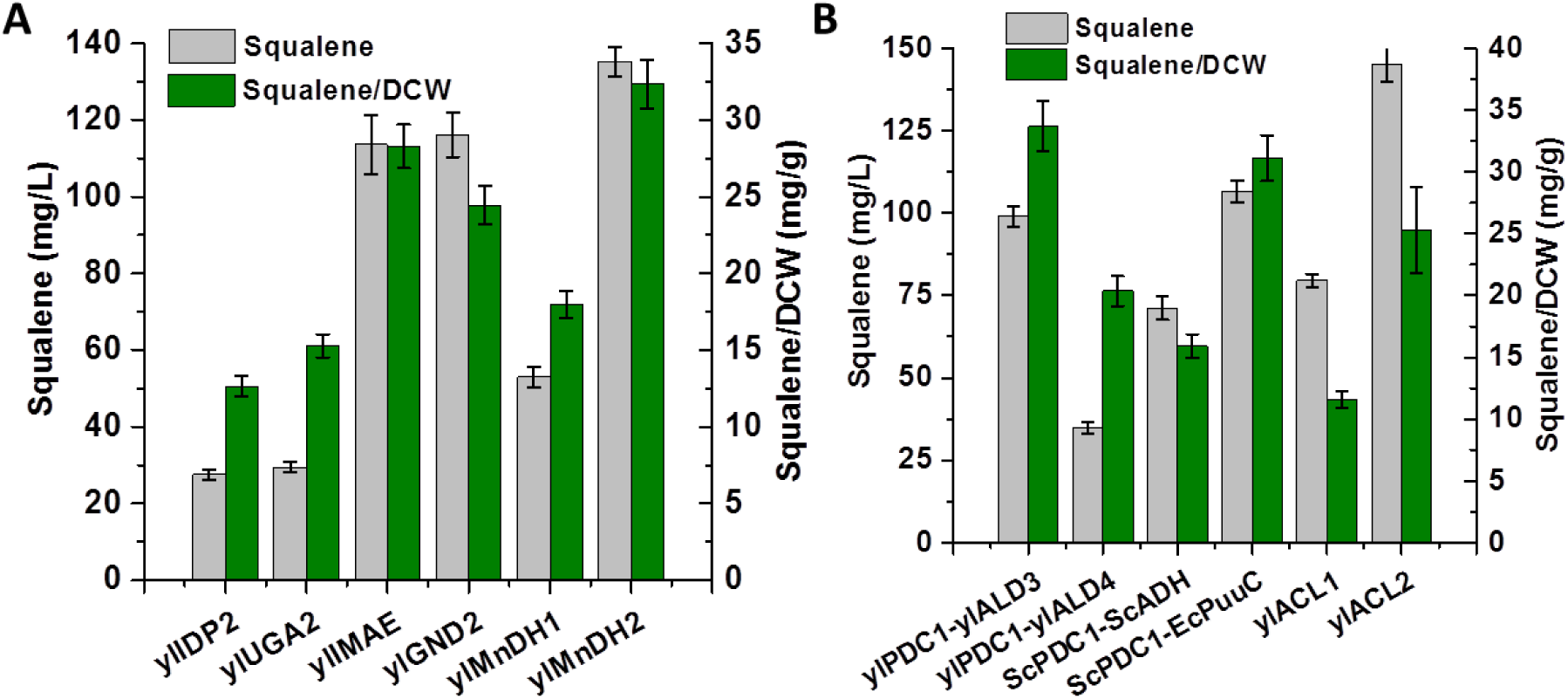
Enhancement of NAPDH and acetyl-CoA precursor pathways to improve squalene production. ScPDC1, pyruvate decarboxylase from *S. cerevisiae*; EcPuuc, aldehyde dehydrogenase from *E. coli*. Other genes are native genes from *Y. lipolytica* and detailed gene annotation could be found in Fig. 1.

Apart from NADPH, acetyl-CoA, is an essential metabolic intermediate connecting glycolysis, Krebs cycle, and glyoxylate shunt pathways. Acetyl-CoA is also the intermediate metabolite participated in lipid synthesis, peroxisomal lipid oxidation and amino acid degradation pathways. It links both anabolism and catabolism, is the starting molecule in MVA pathway. Cytosolic acetyl-CoA was found as a critical precursor to boost secondary metabolite production (Huan Liu, Marsafari, Wang, Deng, & Xu, 2019). For example, engineering alternative cytosolic acetyl-CoA pathways were proven to be efficient strategies to improve fatty acids and isoprenoid production in both Bakers’ yeast and *Y. lipolytica* (Hu Liu, Fan, Wang, Li, & Zhou, 2019; Meadows et al., 2016; van Rossum, Kozak, Pronk, & van Maris, 2016). Therefore, we next investigated whether endogenous and various heterologous acetyl-CoA pathways could improve squalene production. First, we investigated the pyruvate decarboxylase (PDC), acetylaldehyde dehydrogenase (ALD) and acetyl-CoA synthase (ACS) bypass (Fig. 1) and compared the efficiency of this route from *Y. lipolytica*, *S. cerevisiae* and *E.coli* (Fig. 3B). By overexpression of pyruvate decarboxylase (ScPDC) from *S. cerevisiae* and acetylaldehyde dehydrogenase (EcPuuc) from *E.coli*, we obtained only 106.54 mg/L of squalene (Fig. 3B). We observed that the cell growth fitness was negatively impacted due to the expression of heterologous genes, possibly due to the accumulation of the toxic aldehyde intermediate. We next attempted the endogenous ATP citrate lyase, which is the primary acetyl-CoA route to *Y. lipolytica* metabolism. ATP citrate lyase (ACL) was mainly used for supply of the cytosolic acetyl-CoA, which was proven to have two isoforms encoded by two separate genes in *Y. lipolytica* (ACL1 and ACL2) (Nowrousian, Kück, Loser, & Weltring, 2000). Endogenous ylACL1 (YALI0E34793g) and ylACL2 (YALI0D24431g) genes were subsequently tested. A 19.5% increase in squalene synthesis was obtained in the resulting strains *HLYaliS03* with ylACL2 overexpressed along with SQS and ylHMG, leading to the titer of squalene 144.96 mg/L (Fig. 3B). The increase was probably a result of the pushing strategies for acetyl-CoA enrichment by expressing ACL2 so that adequate cytosolic acetyl-CoA could be pushed into the MVA pathway for the synthesis of squalene. Surprisingly, the specific yield reduced to 25.27 mg/g DCW which was caused by the enhancement of cell growth due to the increased lipid content. This increased lipid content may also serve as the storage space to sequestrate squalene in our engineered cell.

### 3.3 Glucose and acetate as media for squalene production

*Y. lipolytica* can grow on a broad range of substrates and convert various organic waste to high-value chemicals (Dobrowolski, Mituła, Rymowicz, & Mirończuk, 2016; Rakicka, Biegalska, Rymowicz, Dobrowolski, & Mirończuk, 2017). For example, it has been reported that *Y. lipolytica* possessed strong acetate utilization pathway that is equivalent or even superior to the hexose utilization pathway, which led to an improvement of triacylglyceride (TAG) production from 100 g/L to 115 g/L in bench-top bioreactors, when the cultivation was switched from glucose media to acetate media (Qiao et al., 2017; J. Xu, Liu, Qiao, Vogg, & Stephanopoulos, 2017). In another work, *Y. lipolytica* was reported to efficiently uptake acetic acid as sole carbon source to produce polyketides up to 4.76 g/L, indicating that acetate may serve as a metabolic “shortcut” to acetyl-CoA with improved carbon conversion efficiency and pathway yield (Huan Liu, Monireh Marsafari, Fang Wang, et al., 2019). In this study, a similar strategy was explored to investigate the conversion process of acetate to squalene by the engineered strain *HLYaliS01*, *HLYaliS02* and *HLYaliS03* (Supplementary Fig. S1). 41 g/L sodium acetate (NaAc), equivalently to 29.5 g/L acetic acid (HAc, 0.5 M) was used to cultivate the engineered strains. *In situ* pH indicator (bromocresol purple) was used to track the pH change and 6 M HCl was used to adjust the pH in the shake flask. Among the engineered strains, the highest squalene titer reached 191.68 mg/L at 168 h in acetate-YNB medium by strain *HLYaliS01*, with 99% of acetic acid depleted and 6.6 g/L biomass produced, yielding of squalene at 29.04 mg/g DCW (Supplementary Fig. S1 B and Table S3). Strain *HLYaliS02* produced 180.28 mg/L squalene at 140 h in acetate-YNB medium with the highest productivity (Supplementary Fig. S1 D). When both engineered strains (*HLYaliS01* and *HLYaliS02*) were cultivated in glucose-YNB medium, 157.81 mg/L and 188.18 mg/L squalene was achieved by strain *HLYaliS01* and *HLYaliS02*, with a yield of 16.53 mg/g DCW and 15.91 mg/g DCW, respectively (Supplementary Fig. S1 A and C, Table S3). This indicates that the mannitol cycle (which is engineered in strain *HLYaliS02* with *SQS-ylHMG-ylMnDH2*) may function well when glucose is used as carbon source. Compared with *HLYaliS01* and *HLYaliS02*, *HLYaliS03* (the strain with *SQS-ylHMG-ylACL2*) produced less squalene in glucose (138.33 mg/L), but similar amount of squalene in acetate (176.8 mg/L) (Supplementary Fig. S1 E and F). The data demonstrated that both glucose and acetate could be utilized as carbon sources to produced squalene by *Y. lipolytica* and acetate as a potential and cheap industrial chemicals had a promising application and commercial value for squalene and terpene production.

To further improve squalene synthesis, we assembled ylACL2 to the plasmid harboring SQS, ylHMG and ylMnDH2. But the engineered strain (*HLYaliS04*) didn’t result in an improved squalene production from neither glucose nor acetate as substrate (Supplementary Fig. S2), possibly due to the metabolic imbalance or gene expression overloading causing burdensome effects to the cell factory.

### 3.4 Shake flask cultivation of engineered strain with pH and carbon/nitrogen ratio optimization

Despite that glucose is the preferred carbon source for cell growth, a similar level of squalene production was detected in our engineered strains (*HLYaliS01* and *HLYaliS02*, *HLYaliS03*). *Y. lipolytica* is a natural lipid producer, accumulating up to 30%~60% cell weight as lipid, which leads to a strong competition for the precursor acetyl-CoA (P. Xu et al., 2017). Meanwhile, cultivation pH and media C/N ratio were two critical factors that affect cellular morphology and growth in *Y. lipolytica* (Szabo, 1999).

In our previous work, we observed a quick declining of cultivation pH from 6 to 3.5 in polyketide synthesis, due to the accumulation of citric acid when glucose was utilized. The pH variations negatively affect strain physiology, alter cell membrane permeability and limit nutrients transportation due to the loss of proton driving force. A significant improvement of polyketide titer was observed by combining PBS buffer with 1 mg/L cerulenin supplementations. Cerulenin is known to irreversibly form a covalent adduct with the active site (cysteine residue) of β-ketoacyl-ACP synthase, inhibiting the elongation of the fatty acid backbone (Huan Liu, Monireh Marsafari, Fang Wang, et al., 2019). Thus, a similar strategy was applied to promote squalene production by strain *HLYaliS02* (shown in Fig. 4 A and Supplementary Fig. S3). When the engineered strain was cultivated in the minimal YNB media with 0.2 M phosphoric buffer solution (PBS, pH 6.0), squalene production was increased to 354.44 mg/L at 168 h (Supplementary Fig. S3), which was an increase of 88.4%, compared with the results from pH uncontrolled (Supplementary Fig. S1 C) experiment. The improvement is ascribed to the better growth fitness under pH control and the biomass of strain *HLYaliS02* reached 13.98 g/L DCW with the squalene specific yield at 25.35 mg/g DCW (Supplementary Table S3). The major byproduct mannitol accumulated up to 2.2 g/L at 48 hour and citric acid reached 7.81 g/L at 96 h; both mannitol and citrate were subsequently reincorporated into cell metabolism (Supplementary Fig. S3). But only ~50% of glucose was utilized in this process which was consistent with our previous work, indicating the supplementation of PO_4_3-buffer may negatively impact the glucose uptake rate. To further enhance squalene synthesis, 1 mg/L cerulenin was supplemented to the minimal YNB-PBS media at 48 h and the squalene production increased to 384.13 mg/L at 188 h, an 8.4% increase compared with the result without cerulenin (Fig. 4 A). A similar fermentation profile of glucose consumption, mannitol, and citric acid accumulation was found: half of glucose was utilized while 14.9 g/L DCW was obtained with the squalene specific yield at 25.78 mg/g DCW (Supplementary Table S3). Byproduct mannitol reached 1.9 g/L at 48 h, but citric acid increased to 9.7 g/L, which was higher than that in the YNB-PBS media without cerulenin supplemented, possibly due to the fact that inhibition of the endogenous fatty acid synthase may prevent citrate from being converted to acetyl-CoA and oxaloacetate by ATP-citrate lyase (encoded by ACL).

**Fig. 4.**
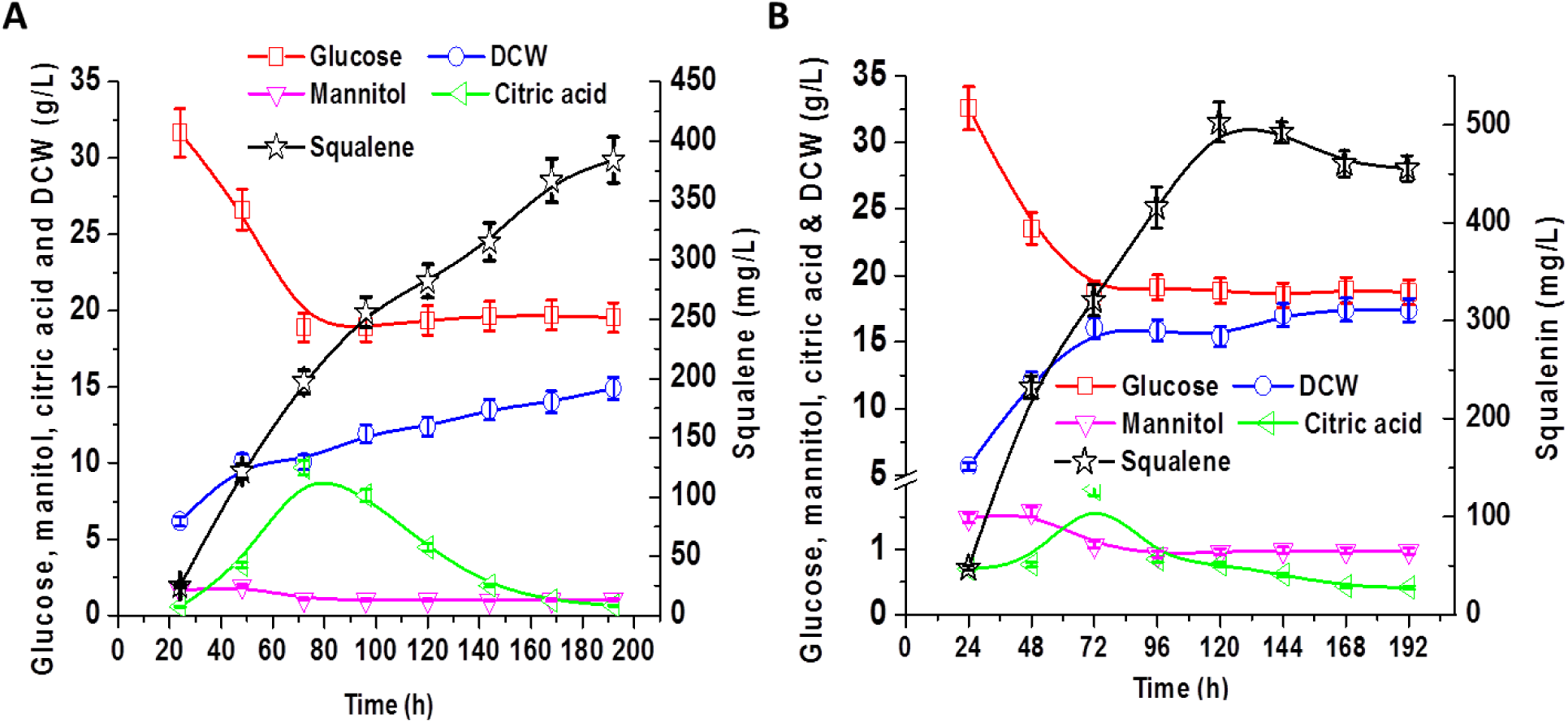
Improving squalene production by controlling pH and C/N ratio with cerulenin supplementation in glucose-minimal media. Fermentation profile of glucose consumption, mannitol, dry cell weight, citric acid and squalene accumulation for strain *HLYaliS02* cultivated in glucose-minimal media conditioned with PBS buffer and supplemented with 1 mg/L cerulenin (A). Fermentation profile of glucose consumption, mannitol, dry cell weight, citric acid and squalene accumulation for strain *HLYaliS02* cultivated in glucose-minimal media conditioned with PBS buffer, supplemented with 1 mg/L cerulenin and C/N ratio 40:1 (B).

We next investigated the effect of C/N ratio on squalene production in YNB-PBS media supplemented with cerulenin (Fig. 4 B). Various C/N ratios including 10:1, 20:1, 40:1, 60:1 and 80:1 was studied (Supplementary Fig. S4). When the C/N ratio was set at 60:1, a similar fermentation profile of glucose consumption, mannitol, dry cell weight, citric acid was obtained, compared to the metabolic profile for C/N 80:1. Squalene titer at C/N 60:1 reached 396 mg/L at 120 h with increased productivity. The highest squalene titer was achieved in the media with C/N ratio 40:1, reaching 502.75 mg/L at 120 h with the yield to 32.6 mg/g DCW (Fig. 4 B, supplementary Table S3), which was 30.8% higher than the squalene production form C/N ratio 80:1 media. We speculated that acetyl-CoA flux was enlarged and flowed to MVA pathway, since we observed less citric acid accumulation (1.9 g/L) at the end of fermentation. However, when the C/N ratio was further reduced to 20:1 or 10:1, adverse effect was obtained with decreasing squalene production (Supplementary Fig. S4). We speculated that the superfluous nitrogen provision may lead more carbons flowing to cell growth. These results illustrated that C/N ratio plays an important role in the redistribution of carbon flux and strongly influenced the accumulation of squalene. Further downregulation of acetyl-CoA carboxylase (ACC) may be required to improve squalene production. ACCase, as the malonyl-CoA source pathway and the acetyl-CoA sink pathway during lipogenesis, was primarily controlled through the phosphorylation of serine residues by Snf1-mediated AMP kinase. Inhibition of fatty acid synthase pathway and nitrogen starvation was proven to be effective strategies to activate Snf1 kinase and slows down ACC1 activity (Seip, Jackson, He, Zhu, & Hong, 2013; Zhang, Galdieri, & Vancura, 2013). It was consistent with our findings that medium C/N ratio was beneficial for squalene synthesis. By applying these engineering strategies, we have obtained an oleaginous yeast strain with a similar squalene level to the strain *S. cerevisiae* (Han, Seo, Song, Lee, & Choi, 2018; Huang et al., 2018). This work highlights the potential of engineering *Y. lipolytica* as a promising microbial platform for efficient synthesis of squalene and terpene-related compounds.

## 4. Conclusion

Squalene is a super lubricant with skin compatibility and thermostability. Traditional source of squalene from shark-hunting or oil plant extraction is cost-prohibitive and not sustainable. Microbial fermentation is considered as a promising route to upgrade sugar renewable feedstock to squalene. By reconstituting mevalonate pathway in yeast, a few of studies have achieved considerable amount of squalene. In this work, we reported the engineering of the oleaginous yeast, *Y. lipolytica*, as a potential host for squalene production. We surveyed a number of HMG-CoA reductase and discovered that endogenous HMG-CoA reductase led to the highest squalene improvement. With the recycling of NADPH from the mannitol cycle, the engineered strain (with MnDH2 overexpression) produced about 180.3 mg/l and 188.2 mg/L squalene from glucose or acetate minimal media, respectively. We identified the optimal NADPH and acetyl-CoA supply mode. Upon overexpression of squalene synthase, HMG-CoA reductase, mannitol dehydrogenase or ATP-citrate lyase, we further optimized the cultivation conditions. The engineered strain fermented with C/N 40:1 media conditioned with PBS buffer with supplementation of 1 mg/L cerulenin produced about 502.7 mg/L squalene in shake flaks. The metabolic byproduct citric acid and mannitol level were also profiled, both byproducts were reincorporated into cell metabolism at the late stage of fermentation. This work may serve as a starting point to harness *Y. lipolytica* as an oleaginous yeast factory for cost-efficient production of squalene or terpene-based chemicals.

## Author contributions

PX conceived the topic and designed the study. HL performed genetic engineering and fermentation experiments. HL and PX wrote the manuscript. PX revised the manuscript.

## Acknowledgments

We would like to acknowledge Bill & Melinda Gates Foundation (grant number OPP1188443) and National Science Foundation (CBET-1805139) for financially supporting this project. The authors would also like to acknowledge the Department of Chemical, Biochemical and Environmental Engineering at University of Maryland Baltimore County for funding support. HL would like to thank the China Scholarship Council for funding support.

