## Supplemental Tables and Figures for "Optimizing mevalonate pathway for squalene production in *Yarrowia lipolytica*"

1. Comparison of squalene production using glucose and acetate by different strains of *Y. lipolytica*

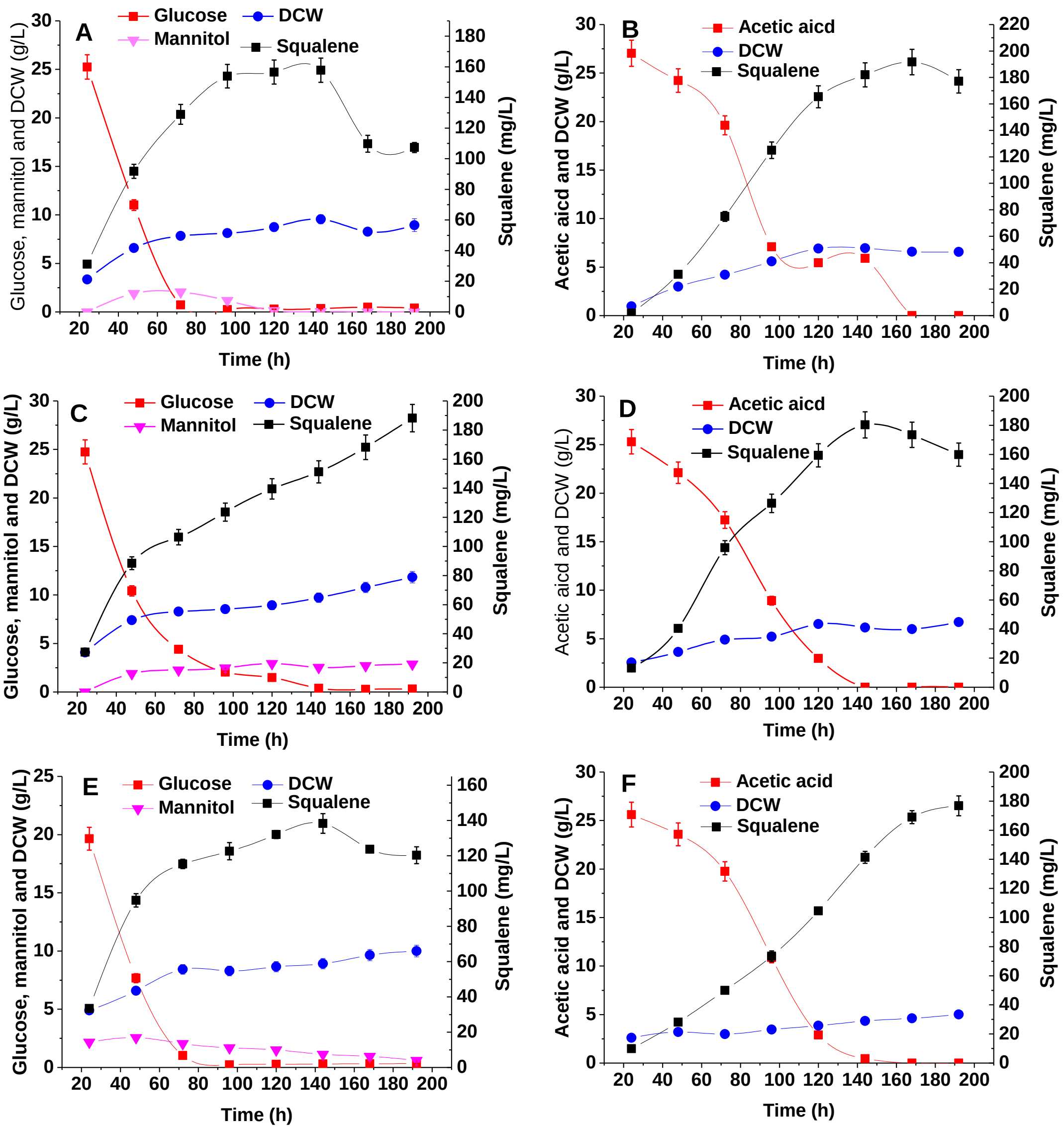


Supplementary Fig. S1 Comparison of squalene production using glucose and acetate as substrate by different engineered strans. Squalene production from minimal media supplemented with glucose and NaAc by strain *HLYaliS01* (A and B); Squalene production from minimal media supplemented with glucose and NaAc by strain *HLYaliS02* (C and D); Squalene production from minimal media supplemented with glucose and NaAc by strain *HLYaliS03* (E and F). When NaAc was used as substrate, the medium pH will increase with the consumption of acetate. We adjusted the pH to 6.0 by adding HCl. Bromocresol purple is a pH-sensitive indicator to track the pH variations of fermentation process.

2. Production of squalene by strain *HLYaliS04* using glucose and acetate as substrate

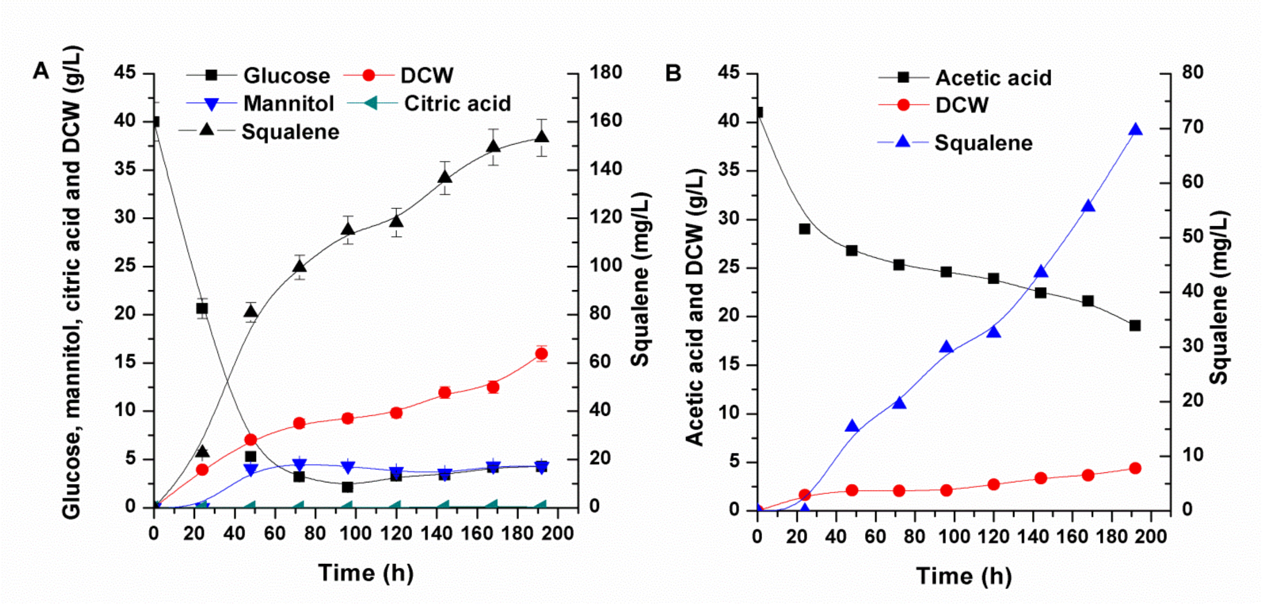


Supplementary Fig. S2 Comparison of squalene production using glucose and acetate as substrate by *HLYaliS04.* Squalene production from minimal media supplemented with glucose by strain *HLYaliS04* (A); Squalene production from minimal media supplemented with NaAc by strain *HLYaliS04* (B).

3. Production of squalene by strain *HLYaliS02* in YNB-PBS media

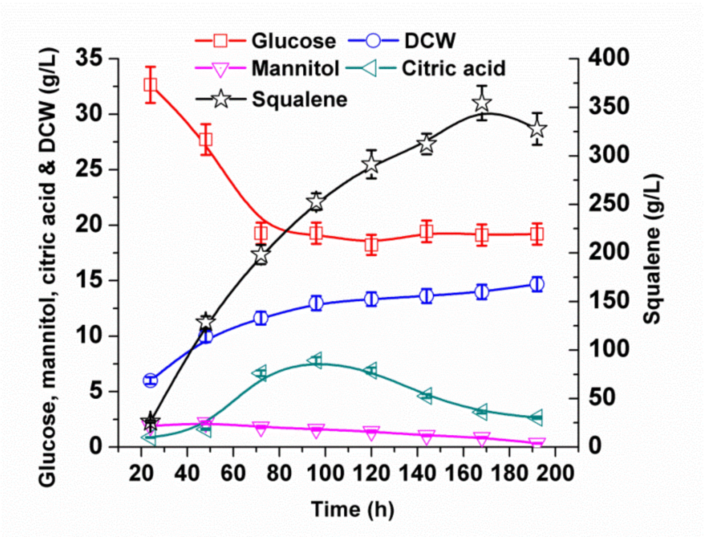


Supplementary Fig. S3 The effect of pH control on the production of squalene by strain *HLYaliS02*. Fermentation profile of glucose consumption, mannitol, dry cell weight, citric acid and squalene accumulation for strain *HLYaliS02* cultivated in glucose-minimal media conditioned with PBS buffer.

4. C/N ratio optimization in squalene production by *HLYaliS02*

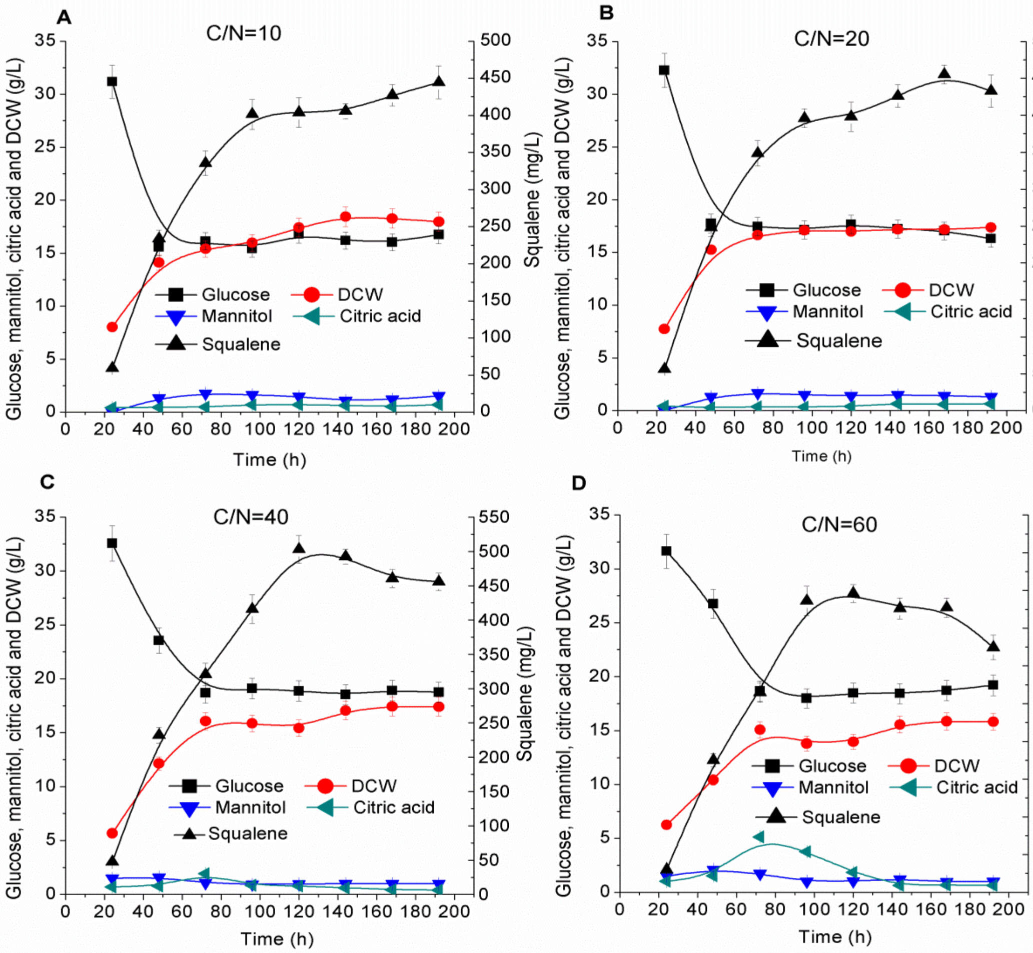


Supplementary Fig. S4 The effect of C/N ratio on the production of squalene by strain *HLYaliS02*. Fermentation profile of glucose consumption, mannitol, dry cell weight, citric acid and squalene accumulation for strain *HLYaliS02* cultivated in glucose-minimal media conditioned with PBS buffer, supplemented with 1 mg/L cerulenin and C/N ratio 10:1 (A), 20:1 (B), 40:1 (C), 60:1 (D).

**Supplementary Table S1. Primers and synthetic oligos/genes used in this study**

| No. | Primer | Nucleotide sequence (5’ >3’) |
|---|---|---|
| 1 | SQS_F | ccgaccagcactttttgcagtactaaccgcagggaaaactcatcgaactgctcttgc |
| 2 | SQS_R | ggggacaggccatggaactagtcggtaccctaatctctcagaggaaacatc |
| 3 | ylHMG_F | ccgaccagcactttttgcagtactaaccgcagctacaagcagctattggaaagattg |
| 4 | ylHMG_R | ggggacaggccatggaactagtcggtaccctatgaccgtatgcaaatattcg |
| 5 | ylt495HMG_F | ccgaccagcactttttgcagtactaaccgcagctacgagaagttgtgcgaaccc |
| 6 | ylt495HMG_R | ggggacaggccatggaactagtcggtaccctatgaccgtatgcaaatattcg |
| 7 | SpHMG_F | ccgaccagcactttttgcagtactaaccgcagacagggaaaaccggccatatag |
| 8 | SpHMG_R | ggggacaggccatggaactagtcggtacctcatgtattttccagaacctgc |
| 9 | SctHMG_F | ccgaccagcactttttgcagtactaaccgcaggaccagcttgttaaaacagaag |
| 10 | SctHMG_R | ggggacaggccatggaactagtcggtaccttatgattttatgcaggttactg |
| 11 | ylErg8_F | ccgaccagcactttttgcagtactaaccgcagaccacctattcggctccggg |
| 12 | ylErg8_R | ggggacaggccatggaactagtcggtaccctacttgaaccccttctcgagc |
| 13 | ylErg10_F | ccgaccagcactttttgcagtactaaccgcagcgactcactctgccccgacttaacg |
| 14 | ylErg10_R | ggggacaggccatggaactagtcggtaccctacttgaaccccttctcgagcc |
| 15 | ylErg12_F | ccgaccagcactttttgcagtactaaccgcaggactacatcatttcggcgccaggc |
| 16 | ylErg12_R | ggggacaggccatggaactagtcggtaccctaatgggtccagggaccgatg |
| 17 | ylErg20_F | ccgaccagcactttttgcagtactaaccgcagtccaaggcgaaattcgaaagcgtg |
| 18 | ylErg20_R | ggggacaggccatggaactagtcggtaccctacttctgtcgcttgtaaatc |
| 19 | ScErg8_F | ccgaccagcactttttgcagtactaaccgcagtcagagttgagagccttcagtgcc |
| 20 | ScErg8_R | ggggacaggccatggaactagtcggtaccttatttatcaagataagtttcc |
| 21 | ScErg12_F | ccgaccagcactttttgcagtactaaccgcagtcattaccgttcttaacttctgc |
| 22 | ScErg12_R | ggggacaggccatggaactagtcggtaccttatgaagtccatggtaaattcg |
| 23 | ScErg20_F | ccgaccagcactttttgcagtactaaccgcaggcttcagaaaaagaaattaggagag |
| 24 | ScErg20_R | ggggacaggccatggaactagtcggtaccctatttgcttctcttgtaaac |
| 25 | ylGPS_F | ccgaccagcactttttgcagtactaaccgcaggattataacagcgcggatttcaagg |
| 26 | ylGPS_R | ggggacaggccatggaactagtcggtacctcactgcgcatcctcaaagtac |
| 27 | ylMAE _F | ccgaccagcactttttgcagtactaaccgcagttacgactacgaaccatgcgaccc |
| 28 | ylMAE _R | ggacaggccatggaactagtcggtaccctagtcgtaatcccgcacatggatg |
| 29 | ylMnDH1 _F | ccgaccagcactttttgcagtactaaccgcagcctgcaccagcaacctacgctactg |
| 30 | ylMnDH1 _R | ggacaggccatggaactagtcggtacctcaaggacaacagtagccgccatc |
| 21 | ylMnDH2 _F | ccgaccagcactttttgcagtactaaccgcagtctggaccttccaccctcgccacg |
| 32 | ylMnDH2 _R | ggacaggccatggaactagtcggtacctcagaggcaaggtagaggtaggtag |

| | | |
|---|---|---|
| 33 | ylIDP2 _F | ccgaccagcactttttgcagtactaaccgcagtccaccaccgctactcgaggcctg |
| 34 | ylIDP2 _R | ggacaggccatggaactagtcggtaccctaagccaggtccttcttcagtc |
| 35 | ylGND2_F | ccgaccagcactttttgcagtactaaccgcagactgacacttcaaacatcaagtg |
| 36 | ylGND2_R | ggacaggccatggaactagtcggtaccttaagcatcgtaagtggaagaag |
| 37 | ylUGA2 _F | ccgaccagcactttttgcagtactaaccgcagttgcgagccctgaataccgtccag |
| 38 | ylUGA2 _R | ggacaggccatggaactagtcggtaccttaaggctgaatgtggggctcgacg |
| 39 | ylPDC1 _F | ccgaccagcactttttgcagtactaaccgcagagcgactccgaaccccaaatggtc |
| 40 | ylPDC1_R | ggacaggccatggaactagtcggtaccctaaacgttggtcttggcagagag |
| 41 | ylALD4 _F | ccgaccagcactttttgcagtactaaccgcagtctcttttcagcaaacttaccctag |
| 42 | ylALD4_R | ggacaggccatggaactagtcggtaccttactgcagcttggcctggtcaaac |
| 43 | ylALD3 _F | ccgaccagcactttttgcagtactaaccgcagcaagttactcttcccgacggaaag |
| 44 | ylALD3_R | ggggacaggccatggaactagtcggtaccctaatccaggttaatgtggac |
| 45 | ylACL1 _F | ccgaccagcactttttgcagtactaaccgcagtctgccaacgagaacatctcccg |
| 46 | ylACL1_R | ggggacaggccatggaactagtcggtaccctatgatcgagtcttggccttgg |
| 47 | ylACL2 _F | ccgaccagcactttttgcagtactaaccgcagtcagcgaaatccattcacgaggc |
| 48 | ylACL2_R | ggggacaggccatggaactagtcggtaccttaaactccgagaggagtgg |
| 49 | ScPDC1_F | ccgaccagcactttttgcagtactaaccgcagtctgaaattactttgggtaaatatttg |
| 50 | ScPDC1_R | ggggacaggccatggaactagtcggtaccttattgcttagcgttggtagc |
| 51 | ScADH_F | ccgaccagcactttttgcagtactaaccgcagtctatcccagaaactcaaaaagg |
| 52 | ScADH_R | ggggacaggccatggaactagtcggtaccttatttagaagtgtcaacaacg |
| 53 | EcPuuc_F | ccgaccagcactttttgcagtactaaccgcagaattttcatcatctggcttactggc |
| 54 | EcPuuc_R | ggggacaggccatggaactagtcggtacctcaggcctccaggcttatccag |

### Supplementary Table S2. Strains and plasmids used in this study

| **Plasmid or strain** | **Relevant properties or genotype** | **Source** |
|---|---|---|
| **Plasmids** | | |
| PYLXP’ | pYaliA1 vector backbone with leucine marker and Ampicillin resistance gene | 1 |
| PYLXP’-SQS | PYLXP’ carrying SQS from *Y. lipolytica* | This study |
| PYLXP’-SQS-ylHMG | PYLXP’ carrying SQS and HMG from *Y. lipolytica* | This study |
| PYLXP’-SQS-ylt495HMG | PYLXP’ carrying SQS and truncated HMG from *Y. lipolytica* | This study |
| PYLXP’-SQS-SctHMG | PYLXP’ carrying SQS from *Y. lipolytica*, truncated HMG from *S. cerevisiae* | This study |

| | | |
|---|---|---|
| PYLXP'-SQS-SpHMG | PYLXP' carrying SQS from *Y. lipolytica*, HMG from *Silicibacter pomeroyi* | This study |
| PYLXP'-SQS-ylHMG-ylErg8 | PYLXP' carrying SQS, Erg8 and HMG from *Y. lipolytica* | This study |
| PYLXP'-SQS-ylHMG-ylErg10 | PYLXP' carrying SQS, Erg10 and HMG from *Y. lipolytica* | This study |
| PYLXP'-SQS-ylHMG-ylErg12 | PYLXP' carrying SQS, Erg12 and HMG from *Y. lipolytica* | This study |
| PYLXP'-SQS-ylHMG-ylErg20 | PYLXP' carrying SQS, Erg20 and HMG from *Y. lipolytica* | This study |
| PYLXP'-SQS-ylHMG-ScErg8 | PYLXP' carrying SQS and HMG from *Y. lipolytica*, Erg8 from *S. cerevisiae* | This study |
| PYLXP'-SQS-ylHMG-ScErg12 | PYLXP' carrying SQS and HMG from *Y. lipolytica*, Erg12 from *S. cerevisiae* | This study |
| PYLXP'-SQS-ylHMG-ylGPS | PYLXP' carrying SQS, GPS and HMG from *Y. lipolytica* | This study |
| PYLXP'-SQS-ylHMG-ylMAE | PYLXP' carrying SQS, MAE and HMG from *Y. lipolytica* | This study |
| PYLXP'-SQS-ylHMG-ylUGA2 | PYLXP' carrying SQS, UGA2 and HMG from *Y. lipolytica* | This study |
| PYLXP'-SQS-ylHMG-ylGND2 | PYLXP' carrying SQS, GND2 and HMG from *Y. lipolytica* | This study |
| PYLXP'-SQS-ylHMG-ylIDP2 | PYLXP' carrying SQS, IDP2 and HMG from *Y. lipolytica* | This study |
| PYLXP'-SQS-ylHMG-ylMnDH1 | PYLXP' carrying SQS, MnDH1 and HMG from *Y. lipolytica* | This study |
| PYLXP'-SQS-ylHMG-ylMnDH2 | PYLXP' carrying SQS, MnDH2 and HMG from *Y. lipolytica* | This study |
| PYLXP'-SQS-ylHMG-ylPDC1-ylALD3 | PYLXP' carrying SQS, PDC1, ylALD3 and HMG from *Y. lipolytica* | This study |
| PYLXP'-SQS-ylHMG-ylPDC1-ylALD4 | PYLXP' carrying SQS, PDC1, ylALD4 and HMG from *Y. lipolytica* | This study |
| PYLXP'-SQS-ylHMG-ScPDC1-ScADH | PYLXP' carrying SQS and HMG from *Y. lipolytica*, PDC1 and ADH from *S. cerevisiae* | This study |
| PYLXP'-SQS-ylHMG-ScPDC1-EcPuuC | PYLXP' carrying SQS and HMG from *Y. lipolytica*, PDC1 from *S. cerevisiae*, PuuC from *E.coli.* | This study |
| PYLXP'-SQS-ylHMG-ylACL1 | PYLXP' carrying SQS, ACL1 and HMG from *Y. lipolytica* | This study |
| PYLXP'-SQS-ylHMG-ylACL2 | PYLXP' carrying SQS, ACL2 and HMG from *Y. lipolytica* | This study |
| **Strains** | | |
| *E. coli NEB5α* | fhuA2 Δ(argF-lacZ)U169 phoA glnV44 Φ80 Δ(lacZ)M15 gyrA96 recA1 relA1 endA1 thi-1 hsdR17 | New England Biolabs |
| *Y. lipolytica po1g* | matA, xpr2-332, axp-2, leu2-270 | Madzak et al. |
| *HLYaliS01* | *Y. lipolytica po1g with vector PYLXP'-SQS-ylHMG* | This study |
| *HLYaliS02* | *Y. lipolytica po1g with vector PYLXP'-SQS-ylHMG-ylMnDH2* | This study |
| *HLYaliS03* | *Y. lipolytica po1g with vecto PYLXP'-SQS-ylHMG-ylACL2* | This study |

| | | |
|---|---|---|
| *HLYaliS04* | *Y. lipolytica po1g with vecto PYLXP'-SQS-ylHMG-ylMnDH2-ylACL2* | This study |

**Supplementary Table S3. Comparison of squalene production among modified strains**

| Strains | Medium | Squalene productivity (mg/L) | DCW (g) | Squalene to DCW yield (mg/g) | Squalene to glucose/acetate yield (mg/g) | Space-time yield (mg/L h) |
|---|---|---|---|---|---|---|
| *HLYaliS01* | Glucose YNB medium | 157.81 | 9.55 | 16.53 | 3.95 | 1.09 |
| *HLYaliS02* | | 188.18 | 11.83 | 15.91 | 4.70 | 0.98 |
| *HLYaliS03* | | 138.33 | 8.91 | 15.53 | 3.46 | 0.96 |
| *HLYaliS04* | | 153.30 | 15.94 | 9.62 | 3.83 | 0.80 |
| *HLYaliS01* | Acetate sodium YNB medium | 191.68 | 6.59 | 29.04 | 6.50 | 1.14 |
| *HLYaliS02* | | 180.28 | 6.16 | 29.27 | 6.11 | 1.25 |
| *HLYaliS03* | | 176.8 | 5.00 | 35.36 | 5.99 | 0.92 |
| *HLYaliS04* | | 69.62 | 4.37 | 15.93 | 2.36 | 0.36 |
| *HLYaliS02* | Glucose YNB medium in PBS buffer | 354.44 | 13.98 | 25.35 | 8.86 | 2.11 |
| *HLYaliS02* | Glucose YNB-PBS medium with cerulenin added | 384.13 | 14.9 | 25.78 | 9.60 | 2.00 |
| *HLYaliS02* | C/N ratio 40:1 glucose YNB-PBS medium with cerulenin added | 502.75 | 15.42 | 32.60 | 12.57 | 4.19 |
